# *Achromobacter* genetic adaptation in cystic fibrosis

**DOI:** 10.1101/2021.01.13.426490

**Authors:** Migle Gabrielaite, Finn C. Nielsen, Helle K. Johansen, Rasmus L. Marvig

**Author notes:** Correspondence to Migle Gabrielaite.

## Abstract

*Achromobacter* is an emerging pathogen in patients with cystic fibrosis (CF) and *Achromobacter* caused infections are associated with more severe disease outcomes and high intrinsic antibiotic resistance. While conventional CF pathogens are studied extensively, little is known about the genetic determinants leading to antibiotic resistance and the genetic adaptation in *Achromobacter* infections.

Here, we analyzed 101 *Achromobacter* genomes from 51 patients with CF isolated during the course of up to 20 years of infection to identify within-host adaptation, mutational signatures, and genetic variation associated with increased antibiotic resistance.

We found that the same regulatory and inorganic ion transport genes were frequently mutated in persisting clone types within and between *Achromobacter* species indicating convergent genetic adaptation. Genome-wide association study (GWAS) of six antibiotic resistance phenotypes revealed the enrichment of associated genes involved in inorganic ion transport genes, transcription gene enrichment in β-lactams, and energy production and translation gene enrichment in the trimethoprim/sulfonamide group.

Overall, we provide insights into the pathogenomics of *Achromobacter* infections in patients with CF airways. Since emerging pathogens are increasingly recognised as an important healthcare issue, our findings on evolution of antibiotic resistance and genetic adaptation can facilitate better understanding of disease progression and how mutational changes have implications for patients with CF.

## Introduction

*Achromobacter* species is an emerging pathogen causing chronic bacterial infections in patients with CF and is a major cause of morbidity and mortality in these patients. (1,2) Analysis of pathogen genomes, i.e. pathogenomics, have shown that within-host pathogen genetic adaptation plays a role in these infections. (3,4) While conventional CF pathogens, e.g. *Pseudomonas aeruginosa* and *Staphylococcus aureus*, are studied extensively, little is known about the extent within- and between-patient genetic adaptation has in *Achromobacter* infections, particularly *A. ruhlandii*, *A. insuavis* and *A. xylosoxidans* as they mainly cause chronic, long-term infections in patients with CF. (5–7) Furthermore, genetic features could be used for successful resistance profile predictions as conventional methods are both time consuming and occasionally do not reflect the *in vivo* susceptibility profiles. (8–10) Knowledge of within-host *Achromobacter* adaptation and genetic factors leading to antibiotic resistance development are key for urgently needed new treatment strategy development and pathogen elimination.

Here, we analyzed 101 previously whole genome sequenced (WGS) *Achromobacter* isolates from 51 patients to investigate the genetic relatedness and within-host genetic changes of *Achromobacter* over the course of up to 20 years of infections. First, we aimed to identify the main gene content differences between- and within-*Achromobacter* isolates. Second, we aimed to identify pace and patterns in genetic changes, and compare it to the other pathogenic bacteria in CF. Finally, we attempted to define the most significant associations between *Achromobacter* genetic features and antibiotic resistance phenotypes. Ultimately, this work on the main *Achromobacter* genomic changes acquired during infections in patients with CF, leads to the possibility of genomic-based disease progression prediction, and improved strategies to track and treat persistent airway infections.

## Materials and Methods

### Bacterial isolates

Our analysis included 101 clinical isolates of *Achromobacter* that were defined in detail previously by Gabrielaite *et al.*, (2020). (11) The isolates were sampled from 51 patients with CF attending the Copenhagen Cystic Fibrosis Center at Rigshospitalet, Copenhagen, Denmark. Over the timespan of 0–20 years (median 6.5 years), 64 isolates from 25 patients were longitudinally collected (median 2 isolates) and 37 isolates from 29 patients were single isolates. 29, 18 and 52 isolates belonged to *A. ruhlandii*, *A. insuavis* and *A. xylosoxidans*, respectively. Furthermore, single isolates belonging to *A. aegrifaciens* and a new genogroup were sequenced.

### Bacterial genome sequencing and definition of clone type

Genomic DNA was prepared from *Achromobacter* isolates using a DNeasy Blood and Tissue kit (Qiagen) and sequenced on an Illumina MiSeq platform, generating 250 base paired-end reads. On average, 1,124,551 reads (range of 350,677–2,118,817) for each of the genomic libraries were obtained. Clone types were defined by Pactyper (12) using the default parameters and species’ core genome defined by GenAPI. (13) Lineage was defined as all isolates belonging to the same species and the same clone type.

### Bacterial genome assembly

Sequence reads from each isolate were corrected and assembled by SPAdes 3.10.1 (14) using default parameters and k-mer sizes ranging from 21 to 127. Assembled contigs were joined to 216 scaffolds on average (92–506).

### Average nucleotide identity calculation

Wrongly annotated public *Achromobacter* species from RefSeq database (15) were identified by calculating ANI with fastANI 1.11 (16) using 95% threshold.

### Aggregated pan-genome generation, characterization and visualization

Aggregated pan-genome was created by clustering all pan-genomes from longitudinally collected lineages and *de novo* assemblies from single-isolate lineages with GenAPI. (13) Every gene in the aggregated pan-genome was then aligned back to the individual pan-genomes/*de novo* assemblies to determine if the gene is (1) non-present in the lineage, (2) present and variable within the lineage or (3) present and non-variable. A matrix for an aggregated pan-genome was generated for 26 longitudinal lineages and 35 single-isolate lineages, and visualized using R (17) with a pheatmap library. (18)

### Bacterial genome alignment and variant calling

Alignments, variant calling and pairwise SNP distance identification for *Achromobacter* isolates were performed using reference genome (GCF_001051055.1 for *A. ruhlandii* (AX01 group), GCF_001558755.2 for *A. insuavis* (AX02 group) and GCF_001457475.1 for *A. xylosoxidans* (AX03 group)) with BacDist (19) workflow that is based on variant calling with Snippy. (20) Sequence alignments on average included 84% (81.72%–89.58%) of the raw sequencing reads for *A. ruhlandii*, 87% (75.45%–92.57%) for *A. insuavis* and 86% (75.95%– 93.61%) for *A. xylosoxidans*. Low-quality variants or variants shared among all isolates were discarded by BacDist.

### Substitution rate estimation

The nucleotide substitution rate (21) estimation was performed for each lineage containing 3 or more isolates sampled at different timepoints (10 lineages in total) by using BEAST 2.6.1. (22) Sequence alignments from BacDist were used as input with the following parameters: (1) sequences were annotated with the sampling date (“dated tips”), (2) HKY substitution model with strict clock parameters, (3) gamma prior for clock rate, (4) prior for population size: 1/X, (5) tree prior: coalescent constant population. MCMC was run for 50,000,000 iterations. Convergence was checked by inspecting an effective sample size and parameter value traces in the Tracer 1.7.1 software. (23) Multiple tests for each sample were performed to ensure reproducibility and convergence. The obtained clock rate (per site per year) was multiplied by the alignment size to obtain a substitution rate per genome per year.

### Virulence and antibiotic resistance gene identification

Orthologs of resistance and virulence genes in 61 *Achrmobacter* lineages were identified with Abricate (24) using VFDB (containing 2,597 genes; retrieved: 2020.09.21) (25) for virulence genes and Resfinder 4 (containing 3,122 genes; retrieved: 2020.09.21) (26) for resistance genes. Gene ortholog was considered present in the corresponding database if the alignment made up minimum 50% of the gene length and its identity was minimum 75%.

### Frequently mutated gene definition

Most frequently mutated genes were defined as the top 1% of all mutated genes for the species. If there were more genes mutated with the same frequency as the 1% most frequently mutated genes, these genes were also included in the final analysis. The identified most mutated genes were annotated by EGGNOG-mapper 2.0.1 (27) using DIAMOND and EGGNOG’s bacterial database.

*P. aeruginosa* orthologs were identified by performing clustering with CD-HIT (28) using word size of 3 and 50% identity thresholds. Joint *Achromobacter* most frequently mutated genes were identified by clustering with CD-HIT (28) with word size of 3 and 80% identity threshold.

### Genome-wide association study with antibiotic resistance phenotypes

DBGWAS 0.5.4 (29) software was used for bacterial genome-wide association analysis using 10 (Amoxicillin-Clavulanate (AMC), Ceftazidime (CAZ), Chloramphenicol (CHL), Colistin (CST), Imipenem (IPM), Meropenem (MEM), Piperacillin-Tazobactam (TZP), Sulfamethizole (SMZ), Tigecycline (TGC) and Trimethoprim-Sulfamethoxazole (SXT)) different antibiotic resistance phenotypes and *de novo* assembled scaffolds of 92 isolates for which the antibiotic susceptibility profiles were available (Gabrielaite *et al.*, 2020 (11) for detailed information on isolate susceptibility). Core genome SNP-based phylogenetic tree was used to correct for population structure while all available annotations of *Achromobacter* genes from UniProt database (30) were used for unitig annotation (271,851 genes; retrieved: 2020.04.19). Ten most significant unitigs for each antibiotic test were used for further analysis as the tool authors advise against using a p-value threshold when testing several phenotypes. (29) *Achromobacter* gene annotations were identified by clustering significant unitig gene sequences with CD-HIT (28) with word size of 3 and 80% identity threshold. Enrichment of COG annotated genes was estimated by comparing the fraction of the associated genes with the fraction in the *Achromobacter* reference genomes for the 5 most frequently associated gene groups where more than one gene was present in a group.

## Results

### *Achromobacter* dataset and incorrect annotations of public genomes

The genomes of 101 *Achromobacter* isolates from the airways of 51 patients with CF attending the Copenhagen Cystic Fibrosis Center at Rigshospitalet were sequenced to follow the within-host evolution and genetic adaptation of the lineages over the initial 0–20 years of infection (Figure 1). All isolates were previously defined as belonging to 5 different species: *A. ruhlandii* (N=29), *A insuavis* (N=18), *A. xylosoxidans* (N=52), *A. aegrifaciens* (N=1) and a new genogroup (N=1). The latter two species were excluded from further analysis as the species contained only a single isolate. The remaining 99 *Achromobacter* genomes were grouped to 61 lineages which were defined as all isolates from one patient belonging to the same species and clone type.

**Figure 1.**
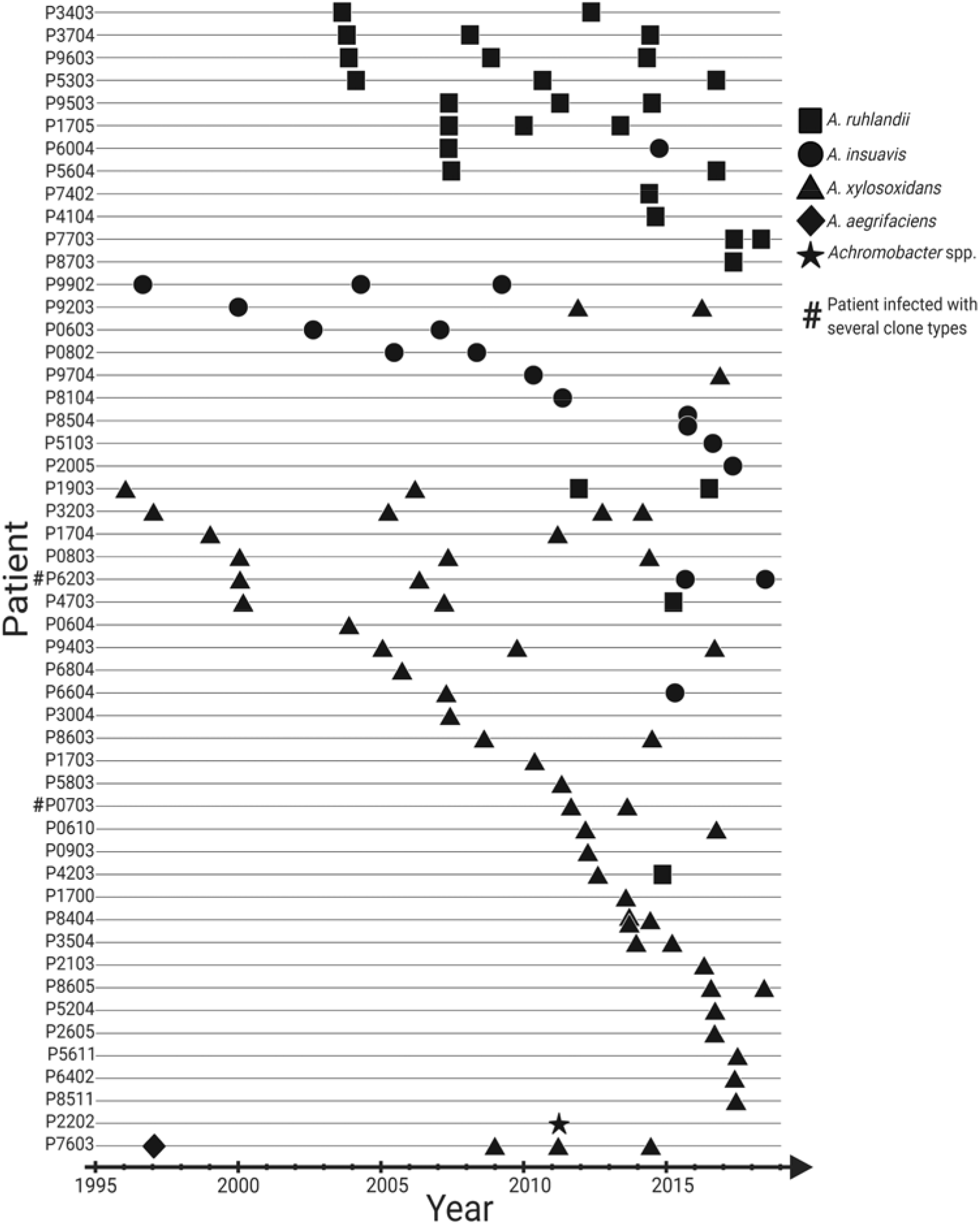
Overview of 101 longitudinally collected *Achromobacter* isolates from patients with CF.

We first performed phylogenetic analysis for all *Achromobacter* genomes available in the RefSeq database (141 samples, Table S1) together with our clinical *Achromobacter* genomes. Our sequenced genomes were widely distributed across the genetic variability observed within *A. insuavis* and *A. xylosoxidans* group; however, *A. ruhlandii* isolates, of which the majority (27/29) belonged to Danish epidemic strain (DES), reflected little genetic variability of *A. ruhlandii* species. Furthermore, our phylogenetic and average nucleotide identity (ANI) analysis revealed that *Achromobacter* annotations are inconsistent among the RefSeq genomes and require corrections to improve species designation (Figure 2A, suggested corrections in Table S1).

**Figure 2.**
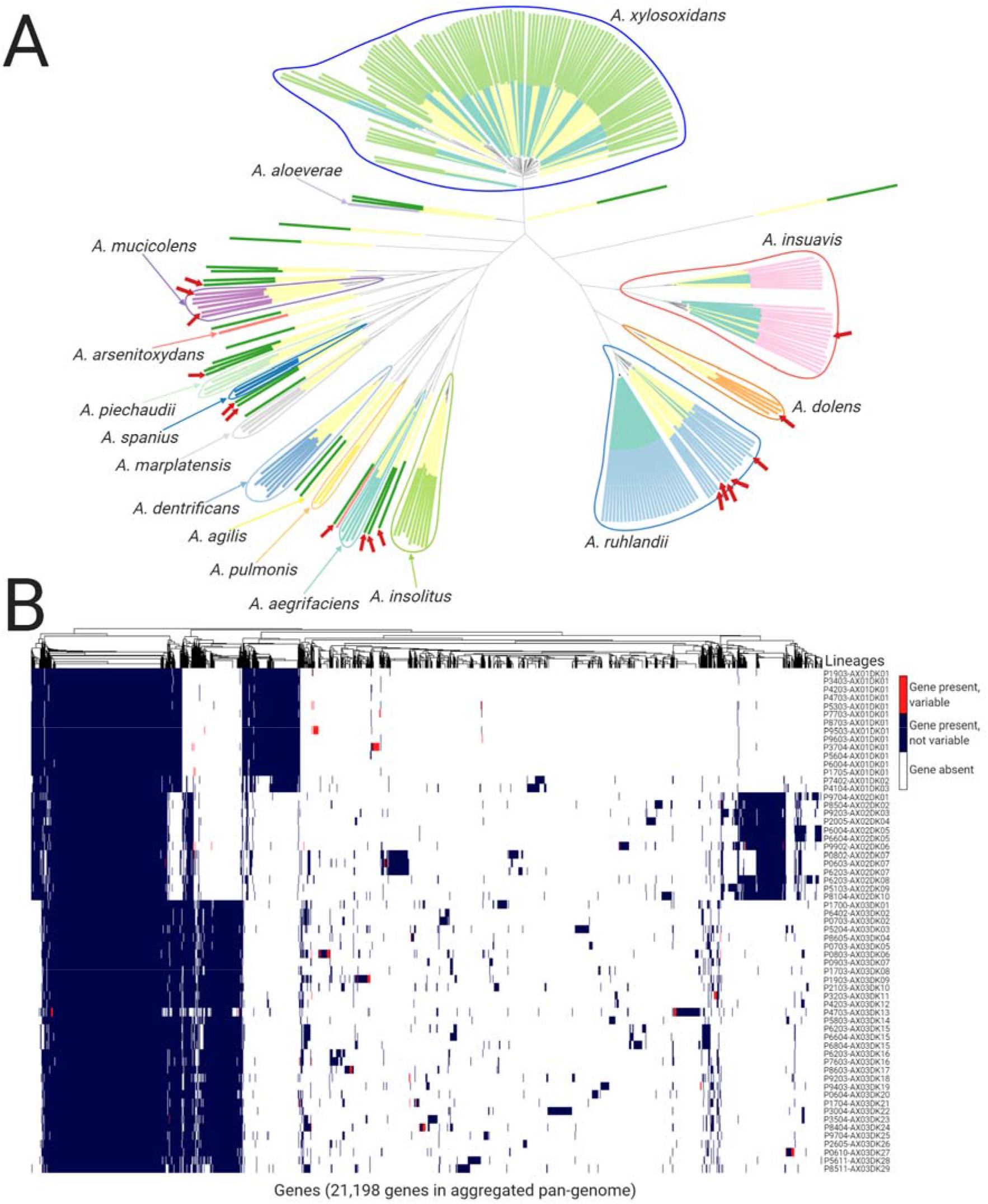
*Achromobacter* genetic differences. (A) Phylogenetic tree based on core genome SNPs of 101 *Achromobacter* isolates from patients with CF (inner layer of colors: turquoise) and 141 *Achromobacter* isolates from RefSeq database (inner layer of colors: yellow). Outer layer of colors corresponds to species annotation with suggested corrections; red arrows mark supposedly incorrect species annotation in RefSeq isolates. The phylogenetic tree can be accessed on Microreact webserver (62). (B) The aggregated pan-genome of 61 *Achromobacter* lineages containing gene presence, absence, and gene variability (i.e., gene is present in some isolates while absent in other isolates within the lineage) information.

### Aggregated pan-genome

Aggregated pan-genome was constructed from pan-genomes of each of the 61 lineages (35 of which were single-isolate lineages). This approach allowed us to account for the nature of the dataset where multiple clonal isolates from the same patient were available. The aggregated *Achromobacter* pan-genome consisted of 21,198 genes: 2,887 core genes, 18,311 accessory genes of which 6,917 genes were unique to a single lineage.

The aggregated pan-genome (Figure 2B) contained *Achromobacter* species-specific genes (649 for *A. ruhlandii*, 648 for *A. insuavis*, and 494 for *A. xylosoxidans*) present in all isolates of the respective species but not in the isolates from other species. Pan-genomes for each *Achromobacter* species were defined by using all bacterial isolates available (18–52) for the species. The size of the species’ pan-genomes contained 7,070–14,833 genes, of which 4,225–5,130 were core genes, 1,940–10,608 accessory genes and 976–3,162 isolate-unique genes (Table 1).

**Table 1.**
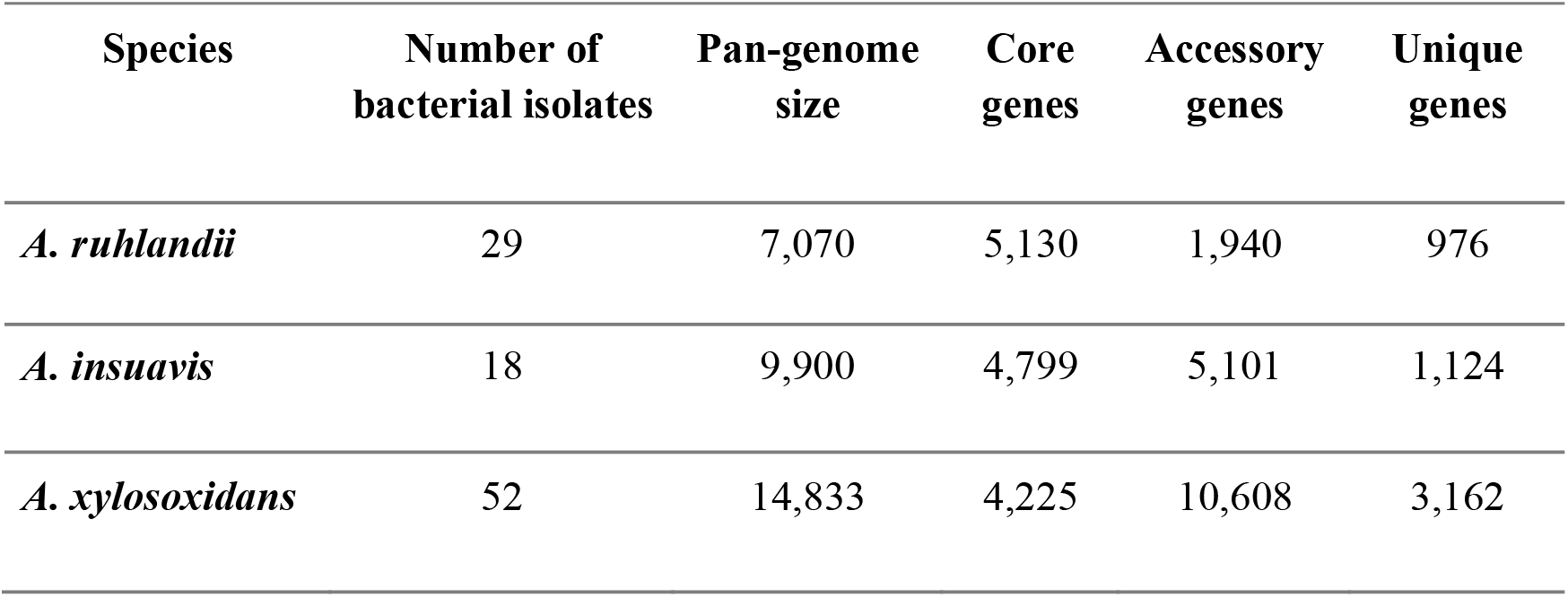
Pan-genome size, number of core, accessory and unique genes for isolates from each Achromobacter species.

| Species | Number of bacterial isolates | Pan-genome size | Core genes | Accessory genes | Unique genes |
| --- | --- | --- | --- | --- | --- |
| *A. ruhlandii* | 29 | 7,070 | 5,130 | 1,940 | 976 |
| *A. insuavis* | 18 | 9,900 | 4,799 | 5,101 | 1,124 |
| *A. xylosoxidans* | 52 | 14,833 | 4,225 | 10,608 | 3,162 |

### *Achromobacter* substitution rates

Within-patient bacterial substitution rate was estimated for lineages where three or more *Achromobacter* isolates from different timepoints were available (five lineages for *A. ruhlandii*, four for *A. xylosoxidans*, and one for *A. insuavis*). The estimated substitution rates for *A. ruhlandii* (DES isolates which are known hypermutators (11,31)) were on average 4.18·10^-6^ (2.71·10^-6^–5.39·10^-6^) SNPs/year/site, 8.77·10^-7^ (6.17·10^-7^–1.13·10^-6^) SNPs/year/site for *A. xylosoxidans* (0332-AX03DK11 hypermutator lineage (11) with the mutation rate of 2.37·10^-6^ SNPs/year/site was excluded), and 1.61·10^-7^

SNPs/year/site for *A. insuavis*. These substitution rates correspond to an average of 21.5, 2.19 and 0.79 SNPs/year/genome for *A. ruhlandii*, *A. xylosoxidans* and *A. insuavis*, respectively (Figure S1).

### Virulence and antibiotic resistance genes carried by *Achromobacter* genomes

From *de novo* assembled genomes of the 99 *Achromobacter* isolates we identified virulence (VFDB database) and antibiotic resistance (Resfinder database) gene orthologs. On average, bacterial isolates carried 2 (0–3) antibiotic resistance gene orthologs and 13 (8–18) virulence gene orthologs. The most frequently carried antibiotic resistance gene was the ortholog of *catB10* which codes for chloramphenicol acetyltransferase. Interestingly, the gene was carried by all *A. ruhlandii* and nearly all (30/33) *A. xylosoxidans* lineages but none of the *A. insuavis* lineages (Figure 3A). Furthermore, OXA-type class D β-lactamase blaOXA-258 was observed in all *A. ruhlandii* isolates and blaOXA-114—in all but one (P7034-AX03DK13) *A. xylosoxidans* lineages. All 13 *A. insuavis* isolates carried one of the following blaOXA genes: blaOXA-243, blaOXA-455, blaOXA-457, blaOXA-458 or blaOXA-459. All blaOXA genes carried by *A. insuavis* had ≥92% nucleotide identity. The latest isolate of DES (P7703-AX01DK01) appears to have acquired a new aph(6)-Id antibiotic resistance gene ortholog encoding for aminoglycoside resistance. (32)

**Figure 3.**
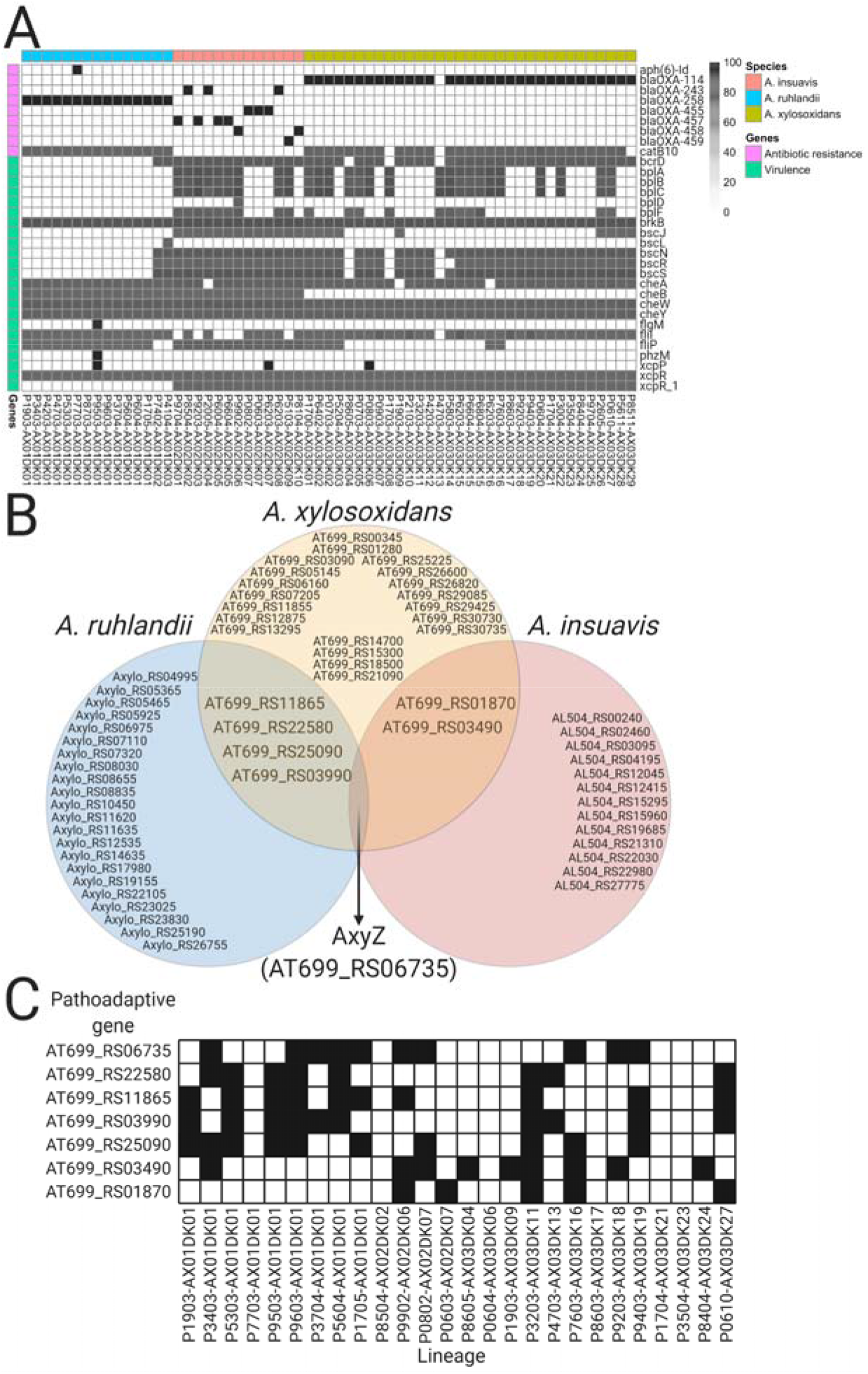
Overview of genetic determinants related to virulence and antibiotic resistance, and pathoadaptive genes. (A) Antibiotic resistance and virulence gene ortholog distribution among lineages. (B) Venn diagram of the most frequently mutated genes and their overlap between the three *Achromobacter* species. (C) Candidate pathoadaptive gene mutation distribution by lineage.

The median number of virulence genes in *A. ruhlandii* DES genomes was markedly lower (N=8) than other *Achromobacter* isolates (N=14; p-value=5.98·10^-8^; Wilcoxon signed-rank test) (Figure 3A). The majority of virulence gene orthologs belonged to the secretion system (N=9), motility (N=7), and endotoxin (N=5) gene orthologs. (25)

### Genetic adaptation: Mutations of the same genes across lineages

To explore within-host genetic adaptation in *Achromobacter*, we first identified the genes which were most frequently mutated within each species. Genes were defined as frequently mutated if they were among the 1% most commonly mutated genes within species. If more than 1% of the genes were mutated with the same frequency, those genes were also included in the analysis. A total of 27, 16, and 28 genes were identified as most frequently mutated for *A. ruhlandii*, *A. insuavis,* and *A. xylosoxidans*, respectively (Figure 3B). The clusters of orthologous groups (COG) functional annotations were performed for all species (Figure 4A; Table S2 for detailed information) with the highest mutation frequency in genes coding for signal transduction (COG T); inorganic ion transport and metabolism (COG P); replication, recombination and repair (COG L); intracellular trafficking, secretion and vesicular transport (COG U); and transcription (COG K).

**Figure 4.**
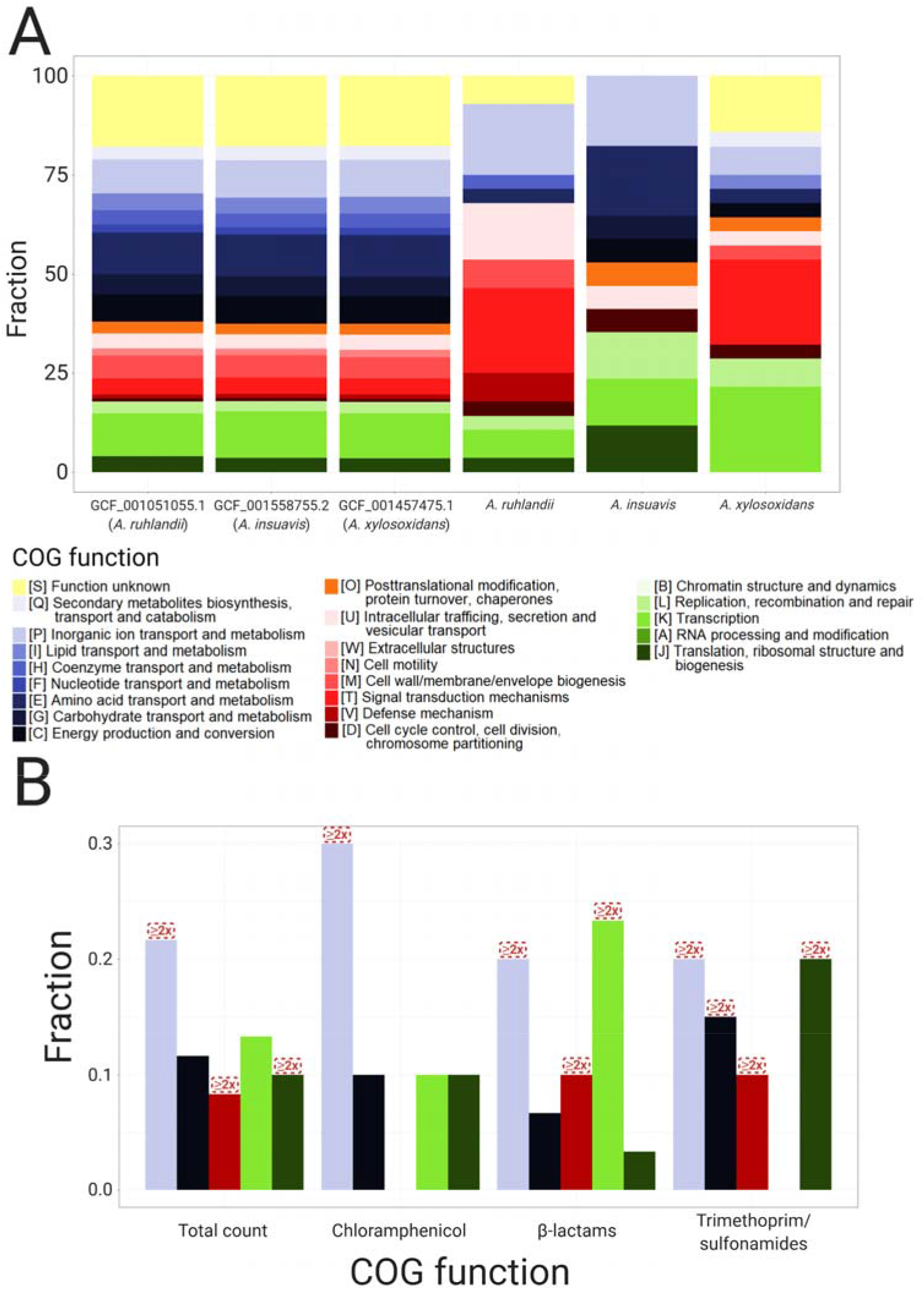
COG annotations of (A) three *Achromobacter* reference genomes and most frequently mutated genes, and (B) frequently associated with antibiotic resistance unitigs. Red boxes mark COGs which are ≥2x enriched in GWAS compared to reference genomes.

Although knowledge is lacking about many *Achromobacter* gene functions, bacterial sequence similarity analysis allowed us to identify possible antibiotic resistance and virulence-related genes among the most frequently mutated genes in *Achromobacter*. After manual literature search 10, 4 and 4 genes were defined as related to antibiotic resistance in *A. ruhlandii*, *A. insuavis* and *A. xylosoxidans*, respectively, whereas 8, 2 and 7 genes were defined as virulence-related genes (Table S2).

Ortholog search of a previously defined list of 52 CF-associated pathoadaptive *P. aeruginosa* genes revealed three orthologs among the most frequently mutated *Achromobacter* genes: *mexZ* (WP_006389199.1), *mexB* (WP_024068614.1) and *gyrA* (WP_049072335.1).

Seven genes were defined as candidate pathoadaptive genes in several *Achromobacter* species. One gene was frequently mutated in all three species (10 out of 26 lineages) and six genes were observed as most frequently mutated in two of the *Achromobacter* species (Figure 3C; Table 2). Six lineages did not acquire a single mutation in any of the seven pathoadaptive genes; however, 13 lineages acquired mutations in three or more genes (Figure 3C). Lineages which acquired mutations in three or more pathoadaptive genes acquired significantly less non-synonymous mutations than lineages with less mutated pathoadaptive genes (p-value=4.99·10^-2^; Fisher’s exact test). Furthermore, lineages only acquired non-synonymous mutations in the seven candidate pathoadaptive genes (Table S3).

**Table 2.** Seven most frequently mutated genes and their function.

| RefSeq ID<br>(Locus tag) | No. of<br>species | No. of<br>lineages | Product | Function |
| --- | --- | --- | --- | --- |
| WP_006389199.1<br>(AT699_RS06735) | 3 | 10 | DNA-binding transcriptional<br>regulator AxyZ | Antibiotic<br>resistance |
| WP_006227290.1<br>(AT699_RS01870) | 2 | 5 | NAD(P)/FAD-dependent<br>oxidoreductase | Metabolic<br>pathways |
| WP_013391523.1<br>(AT699_RS03490) | 2 | 10 | penicillin-binding protein 2 | Antibiotic<br>resistance |
| WP_006392856.1<br>(AT699_RS03490) | 2 | 9 | ABC transporter substrate-<br>binding protein | Transport |
| WP_006394390.1<br>(AT699_RS11865) | 2 | 9 | multidrug efflux RND<br>transporter permease | Antibiotic<br>resistance |
| WP_006387572.1<br>(AT699_RS22580) | 2 | 8 | Signal transduction histidine<br>kinase | Two-component<br>signaling |
| WP_006221301.1<br>(AT699_RS25090) | 2 | 10 | TonB-dependent hemin,<br>ferrichrome receptor | Transport |

The only pathoadaptive gene which was frequently mutated among all three species—*axyZ*— encodes TetR family transcriptional regulator of the RND-type efflux system and is associated with innate *Achromobacter* antibiotic resistance. (33,34) Furthermore, we observed higher overall antibiotic tolerance by isolates which acquired mutations in *axyZ* gene with highest antibiotic resistance increase against piperacillin/tazobactam (p-value=3.28·10^-2^; Fisher’s exact test) and meropenem (p-value=2.87·10^-3^; Fisher’s exact test).

The ratio of non-synonymous to synonymous substitutions (dN/dS) was significantly different between the 1% most frequently mutated genes and non-frequently mutated genes in *Achromobacter* (dN/dS99%=1.21 vs dN/dS1%=2.10; Fisher’s exact test; p=3.4·10^-4^, respectively).

Finally, we investigated gene loss and acquisition patterns in 26 longitudinally collected *Achromobacter* lineages. We observed genes to be 2 times more often lost than acquired, and lost/acquired in groups rather than individually; however, no convergent evolution patterns in *Achromobacter* gene loss/acquisition were identified (detailed analysis in Text S1).

### Genome-wide association between *Achromobacter* genotypes and antibiotic resistance

To test for associations between bacterial genetics and antibiotic resistance phenotypes we performed unitig-based DBGWAS analysis. Out of 21 antibiotics where the resistance was phenotypically tested, only 10 antibiotics had both susceptible and resistant isolates from all three *Achromobacter* species. Accordingly, we performed the association analysis for these 10 antibiotics (see Materials and Methods for detailed information). Unitigs passed a 5% FDR corrected q-value threshold only for CHL, IPM, MEM, TZP, SMZ and SXT. Ten most significant unitigs were used for the six remainder association test analysis, resulting in 60 genes (50 unique genes) significantly associated with antibiotic resistance phenotypes (Figure 4B; Table S5). The most abundant group of associated unitigs belonged to inorganic ion transport (N=13; 2.4x enriched; COG P) genes. The other four most abundantly associated genes belonged to transcription (N=8; COG K); energy production and conversion (N=7; COG C); translation, ribosomal structure and biogenesis (N=6; COG J), and defense mechanism (N=5; COG V) groups. Furthermore, transcription genes were enriched in the β-lactam (IPM, MEM and TZP) antibiotic group (N=7; 2.1x enriched) while translation, ribosomal structure and biogenesis genes were 5.4x enriched in trimethoprim/sulfonamides (SMZ and SXT) group (N=4). Defense mechanism genes were 9.2x and 9.0x enriched in β-lactam and trimethoprim/sulfonamide groups, respectively while energy production and conversion genes were 2.9x enriched in a trimethoprim/sulfonamide group.

Of 9 antibiotic resistance genes from the ResFinder 4 database which were present in the aggregated pan-genome, none were associated with antibiotic resistance phenotypes in the GWAS analysis. Furthermore, core and accessory genes were equally associated with antibiotic resistance phenotypes (Table S5).

## Discussion

*Achromobacter* is an emerging pathogen causing chronic respiratory tract airway infections in patients with CF; however, the genetic epidemiology of these infections is little understood. We sequenced and analyzed 101 genomes of *Achromobacter* isolates from 51 patients with CF which is the largest longitudinally collected *Achromobacter* genome dataset available to-date. This allowed us to investigate the population genomics and within-host adaptation, including genome-wide association analysis with antibiotic resistance phenotypes.

Phylogenetic analysis of our dataset with 141 publicly available *Achromobacter* genomes from the RefSeq database revealed that our dataset well represented the genetic diversity of *A. xylosoxidans* and *A. insuavis* species. However, because of DES overrepresentation, *A. ruhlandii* isolates did not reflect the species genetic diversity. Furthermore, ANI together with core-genome based phylogenetic analysis revealed that more than 10% of publicly available *Achromobacter* genomes are supposedly misannotated in the RefSeq database which we anticipated to unravel to ease future research on *Achromobacter* (Table S1).

From the aggregated pan-genome analysis we showed that core genome sizes were comparable between *A. ruhlandii*, *A. insuavis* and *A. xylosoxidans* which are similar to core genome size identified previously by Li *et al.* 2013 (35). Nevertheless, the number of accessory and unique genes in species’ pan-genomes varied greatly as a result of the different number of independent genomes for each species. *A. ruhlandii* isolates mostly belonged to DES, which is spread through patient-to-patient transmission (11,31), therefore, have lower pan-genome plasticity than *A. xylosoxidans* or *A. insuavis*.

Unlike many bacterial species, substitution rates for *Achromobacter* are not known. (36) Our *A. ruhlandii* dataset only consisted of hypermutator DES lineages which led to a high substitution rate estimate, and our substitution rate estimate for *A. insuavis* was based on only one longitudinally sampled lineage. *A. xylosoxidans* (8.77·10^-7^ SNPs/year per site) substitution rate is comparable to other Gram-negative bacterial species: *P. aeruginosa* (4.0·10^-7^ SNPs/year per site) (37), *Shigella sonnei* (6.0·10^-7^ SNPs/year per site) (38), *Echerichia coli* (2.26·10^-7^ SNPs/year per site) (39). The substitution rate is substantially lower than *S. aureus* (1.87·10^-6^ SNPs/year per site) (40) or *Klebsiella pneumoniae* (1.9·10^-6^ SNPs/year per site) (41).

While it was previously suggested to use blaOXA genes for *Achromobacter* species typing (42–44), we showed that in some cases such strategy would not be sufficient for species identification as isolates can carry none of the blaOXA genes. Furthermore, we identified that *A. insuavis* can carry one out of several highly similar blaOXA genes which might further complicate such species typing approach. We also show that none of the *A. insuavis* isolates carried *catB10* chloramphenicol resistance gene ortholog; however, these identified resistance genes alone were not sufficient to explain the differences in antibiotic resistance phenotype between lineages and species.

Moreover, virulence gene ortholog analysis revealed that *Achromobacter* carries several virulence factors with markedly less virulence genes in *A. ruhlandii* DES strain which further supports the adaptive trade-off evolution hypothesis that virulence genes are not required or are selected against in chronic infections. (36) However, several virulence gene orthologs coding for host cell invasion (*cheW* and *cheY*) and facilitating evasion of the host immune response (*brkB* (45)) were observed in all lineages. Furthermore, our findings of secretion system, in particular type III secretion system, gene orthologs as the most prevalent virulence genes in *Achromobacter* are in line with findings by Jeukens *et al.* 2017 and Li *et al.* 2013. (35,46)

Among the candidate pathoadaptive genes (frequently mutated genes), we identified multiple antibiotic resistance genes which were markedly more frequently mutated among *A. ruhlandii* isolates than *A. insuavis* or *A. xylosoxidans* isolates. This phenomenon might signal about the continuous adaptive evolution even in highly antibiotic-resistant strains such as DES. (1) Nonetheless, more antibiotic resistance and virulence genes among frequently mutated genes might be identified if gene annotation of *Achromobacter* reference genomes improved. Overall, our identified candidate pathoadaptive genes (belonging to signal transduction; inorganic ion transport and metabolism; intracellular trafficking; transcription; and replication gene functional classes) are comparable to the observations in *P. aeruginosa* infecting patients with CF (47) and other smaller-scale studies on *Achromobacter*. (46) Furthermore, we highlight that not all *Achromobacter* lineages seem to undergo the same amount of selective pressure, and we show from the 7 pathoadaptive *Achromobacter* gene analysis that lineages tended to either have acquired not more than 2 pathoadaptive mutations and be under stronger positive selection or have acquired mutations in more pathoadaptive genes and be under weaker/neutral selection. These findings are comparable to findings from genetic adaptation studies in *P. aeruginosa* from patients with CF airway. (47) Overall, significantly higher dN/dS ratio in the 27, 16 and 28 most frequently mutated *A. ruhlandii*, *A. insuavis* and *A. xylosoxidans* genes confirms that there is strong selective pressure for changes in these genes during adaptation to the patients with CF airway. Altogether, dN/dS>1 in both frequently and non-frequently mutated genes show that more than the top 1% mutated genes are under selective pressure; nonetheless, a larger dataset is needed to identify more genes without sacrificing analysis accuracy.

*AxyZ* (*mexZ* ortholog), which was the only candidate pathoadaptive gene in all three species, is involved in the development of multidrug resistance by regulating AxyXY-OprZ RND-type efflux system, hence is crucial during adaptation to the host environment. (34) *AxyZ* ortholog *mexZ* in *P. aeruginosa* is established among pathoadaptive genes directly associated with increase in antibiotic resistance. (48–50) Furthermore, mutations in *axyZ* could partially explain the increased tolerance to piperacillin/tazobactam and meropenem.

The observed gene loss and acquisition patterns were comparable to the ones observed in *P. aeruginosa*; however, unlike in *P. aeruginosa*, no convergent loss or acquisition of gene clusters was observed. (51)

To further explore the differences in antibiotic susceptibility between *Achromobacter* isolates, we performed a k-mer based GWAS analysis. Limited number of bacterial isolates and high innate resistance to certain antibiotics restricted our analysis to only six successful GWAS associations which revealed that inorganic ion transport genes contribute to antibiotic resistance development in all six antibiotics. Changes in inorganic ion transport genes (COG P) could have a secondary influence on antibiotic resistance as such changes help overcome the problem of iron deficiency in human airways allowing better intrinsically resistant bacteria survival despite the presence of antibiotics. Similar patterns were previously identified in other bacteria causing chronic infections in patients with CF. (36,52,53) Another possible explanation is that many inorganic ion transport genes are related to efflux pumps and transporter genes which markedly contribute to increase in antibiotic resistance. (54,55) The observed enrichment of transcription genes (COG K) associated with resistance to β-lactams could be explained by changes in transcriptional regulation of intrinsic antibiotic resistance, efflux pump and cell wall protein coding genes. (54,56,57) Enrichment of translation and ribosomal structure genes (COG J), and energy production genes (COG C) in trimethoprim/sulfonamide group might be due to altered metabolism and changes in energy production leading to bacterial persistence and escape of antibiotic effect. (58,59) Ultimately, GWAS is a promising approach for a systematic innately complex bacterial resistance analysis which could be applied to better understand the genetics of antibiotic resistance development.

Our study has several limitations. First, even larger studies are necessary to further characterize and identify the genetic adaptation of *Achromobacter* during CF airway infections. Second, the lack of genome annotation and overall knowledge about *Achromobacter* limited the interpretation of putative pathoadaptive genes and genes associated with resistance phenotypes. Finally, a single isolate at a given time point is not sufficient to completely reflect the genetic diversity of the bacterial population (60,61); therefore, some of our findings might be the result of diversification and not the fixation of the adaptive mutations in *Achromobacter.*

## Conclusions

In conclusion, by using the largest dataset to-date of *Achromobacter* clinical isolates from patients with CF, we used a comprehensive analytical framework for thorough bacterial genomic data analysis. Thus, we identified pathoadaptive and antibiotic resistance genes in *Achromobacter* causing CF airway infections. Furthermore, we showed that current knowledge about antibiotic resistance gene presence or mutations in those genes cannot sufficiently explain the resistance phenotypes, and GWAS offers a new approach of addressing this problem. The gained knowledge allows us to better understand the requirements for successful *Achromobacter* adaptation during infection in airways of patients with CF which could help accurately predict antibiotic susceptibility and clinical progression of *Achromobacter* infections, and further the development of urgently needed optimized treatment strategies.

## List of abbreviations

AMC: Amoxicillin-Clavulanate
ANI: Average nucleotide identity
CAZ: Ceftazidime
CHL: Chloramphenicol
COG: Clusters of Orthologous Groups
CST: Colistin
CF: Cystic fibrosis
DES: Danish epidemic strain
GWAS: Genome-wide association study
IPM: Imipenem
MCMC: Markov chain Monte Carlo
MEM: Meropenem
TZP: Piperacillin-Tazobactam
SMZ: Sulfamethizole
SNP: Single nucleotide polymorphism
TGC: Tigecycline
SXT: Trimethoprim-Sulfamethoxazole
WGS: Whole genome sequence

## Availability of data and materials

*Achromobacter* whole genome sequencing data is available at European Nucleotide Archive under study accession number PRJEB39108.

## Acknowledgments

Ulla Johansen is thanked for expert technical assistance and Niels Høiby is thanked for collecting the earliest *Achromobacter* isolates. All figures were partly or completely created using BioRender (https://biorender.com/).

## Contributions

R.L.M. and H.K.J conceived the study. R.L.M., H.K.J. and F.C.N supervised the study. M.G. and R.L.M. designed the bioinformatics workflows for the analysis. M.G. conducted the analysis. M.G., and R.L.M. analyzed and interpreted the data. M.G. prepared the manuscript draft and visualizations. R.L.M., H.K.J., and F.C.N reviewed and edited the draft.

## Ethics declarations

Use of the stored clinical isolates was approved by the local ethics committee at the Capital Region of Denmark RegionH (registration number H-4-2015-FSP).

## Supplementary information

**Text S1.** *Achromobacter* gene presence-absence analysis results.

**Table S1.** Table of publicly available RefSeq *Achromobacter* sequences with original and new proposed annotations which were based on ANI and core genome SNP-based phylogeny.

**Table S2.** List of most frequently mutated genes in *Achromobacter ruhlandii, Achromobacter insuavis* and *Achromobacter xylosoxidans* with their COG function, corresponding E-value, gene description and gene relatedness to antibiotic resistance or virulence with references defining gene’s relatedness to antibiotic resistance or virulence are provided where relevant.

**Table S3.** All mutations acquired by lineages in the 7 pathoadaptive genes which were frequently mutated in at least two *Achromobacter* species. Amino acid coordinates are based on the corresponding species’ reference genome genes.

**Table S4.** Table of pan-genome, core, accessory and unique genes in *Achromobacter* lineages. Variable genes are separated by whether they were (1) lost or acquired and (2) variable in a group or individually.

**Table S5.** The most significant GWAS associations (q-value<0.05) and their annotations.

**Figure S1.** Substitution rate per year average with 95% confidence intervals for *Achromobacter ruhlandii* (DES) and *Achromobacter xylosoxidans*.

**Figure S2.** COG annotation of the aggregated pan-genome and most frequently lost or acquired genes.

