## Supplementary material for "*Achromobacter* genetic adaptation in cystic fibrosis": Table S1

As a crucial part of bacterial adaptation and evolution, we investigated the most frequently lost and acquired genes in the 26 longitudinally sampled *Achromobacter* lineages. First, *de novo* assembled genomes were annotated using Prokka version 1.12 [[1]](http://sciwheel.com/work/citation?ids=429229&pre=&suf=&sa=0) with the settings of a minimum contig length of 200 nucleotides and using a manually created annotation database for *Achromobacter* (GCF_001051055.1 (*A. ruhlandii*), GCF_001558755.2 (*A. insuavis*) and GCF_001457475.1) (*A. xylosoxidans*). Further analysis was based on longitudinal lineages by GenAPI software with default settings. [[2]](http://sciwheel.com/work/citation?ids=9355772&pre=&suf=&sa=0) Genes shorter than 150 nucleotides were excluded from further analysis by default. We defined a gene as lost if it was present in the first isolate but absent in one or more of the later isolates and a gene was defined as acquired if it was absent in the first isolate but present in one or more of the later isolates. The lineage pan-genomes contained on average 5,940 (5,709–6,377) genes of which on average 62 (14–183) were lost or acquired.  In total, we found 1,464 genes to be variable within lineages and 735 of these were unique to a single lineage in the aggregated pan-genome (Figure 2B). We observed that genes were 2 times more often lost than acquired and lost/acquired in groups 37 times more commonly than individually (Table S4).

Most frequently lost or acquired genes were defined if a gene was lost/acquired in a minimum of 3 lineages. In total, 38 genes passed these criteria; however, after the manual inspection, one gene was defined as a false positive: the gene was manually assessed as present, yet the gene was not assembled with a minimum requirement of 25% of the gene length coverage, thus was predicted to be absent by GenAPI. The remaining 37 genes that were frequently lost or acquired, were annotated with EGGNOG-mapper: transport and metabolism genes were more frequently lost/acquired, especially amino acid transport and metabolism genes, when compared to the composition of the aggregated pan-genome. Contrary, structural, cellular organization and cell cycle genes; reproduction; and gene expression regulating genes were less frequently lost/acquired (Figure S2).
