## Supplementary material for "*Achromobacter* genetic adaptation in cystic fibrosis": Figure S1

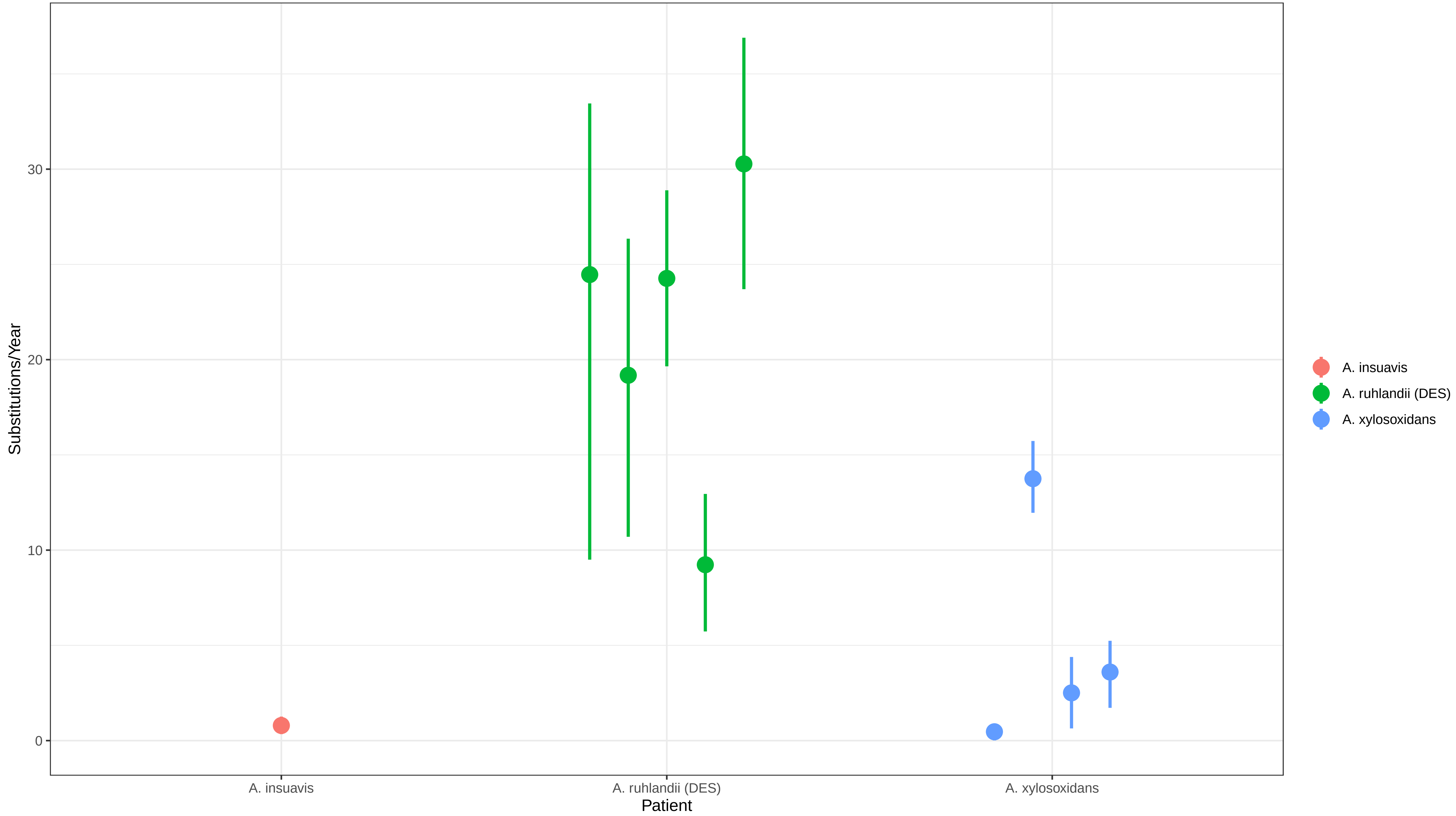


The tree height for the A. insuavis model is large, thus the estimated substitution rate might be too low
