## Supplementary figures and images for "*Achromobacter* genetic adaptation in cystic fibrosis"

### Figure S2

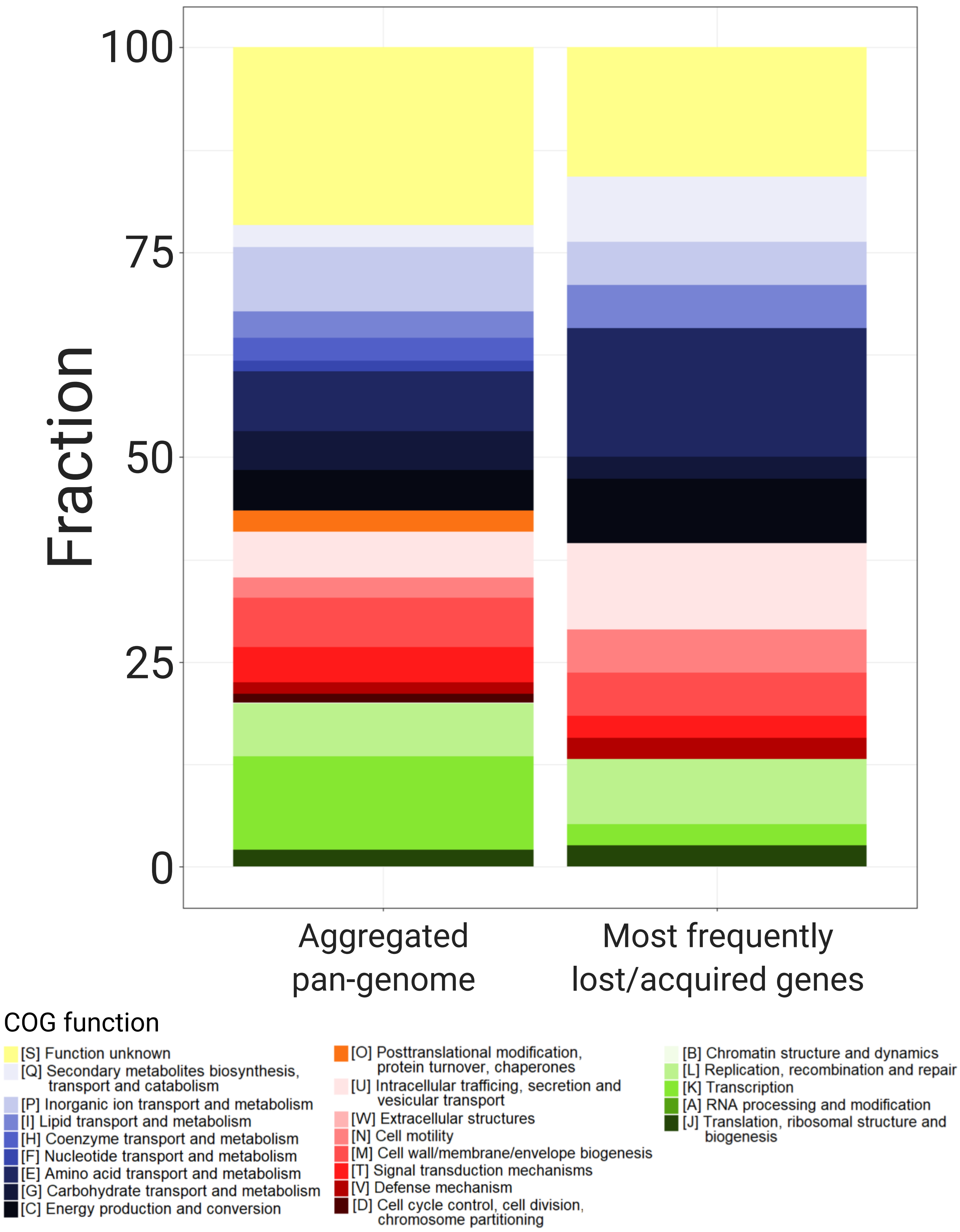
